# Neuron-derived extracellular vesicles in plasma present a potential non-invasive biomarker for Huntingtin protein and RNA assessment in Huntington disease

**DOI:** 10.1101/2025.07.17.665403

**Authors:** Inês Caldeira Brás, Yuanyun Xie, Amber Lee Southwell

## Abstract

Huntington disease (HD) is a neurodegenerative disease caused by a trinucleotide repeat expansion in the HTT gene encoding an elongated polyglutamine tract in the huntingtin (HTT) protein. The use of biomarkers has become a major component in preclinical studies focusing on HTT lowering strategies. Quantification of soluble mutant HTT (mHTT) in cerebrospinal fluid (CSF) has served as a pharmacodynamic readout and as potential disease progression biomarker. However, development of future assays for HTT measurement from other biofluids, such as blood, will facilitate the access to human samples since CSF collection is an invasive outpatient procedure. Brain cells, in particular neurons, secrete extracellular vesicles (EVs) that cross the blood-brain barrier and circulate in blood. Importantly, EVs have been identified to be involved in HTT export from cells to the extracellular space. However, it is unknow which vesicle subtype correlates better with HD progression. Our work investigates the potential of EVs as non-invasive sources of clinical biomarkers in liquid biopsies.

We developed an optimized ultracentrifugation protocol for the purification of ectosomes and exosomes from human samples and plasma of humanized HD mouse models. Ectosomes are larger vesicles that bud from the plasma membrane of cells, whereas exosomes originate from multivesicular bodies and are afterwards released to the extracellular space. Consistent with previous published data in other model systems, ectosomes isolated from plasma of the Hu97/18 mouse model contain both wild-type (wt) and mHTT in higher levels than in exosomes. Similar results were observed in media from HD induced pluripotent stem cells (iPSCs)-differentiated neurons and in Hu97/18 primary neuronal cultures. Interestingly, we also found higher levels of HTT transcripts in this EV subtype. We further demonstrate that initial storage of the samples using a slow freezing protocol preserves HTT and EV protein marker levels, highlighting the importance of sample preparation for EV isolation and analysis. Our results also show that plasma contains vesicles originated from neuronal cells that can be isolated using neuron-specific markers, such as ATPase Na+/K+ transporting subunit alpha 3 (ATP1A3), allowing the evaluation of HTT levels in the brain through vesicles circulating in the blood.

Overall, our results demonstrate that HTT protein measurement from EVs isolated from blood can be a potential less-invasive disease biomarker. We also demonstrate that EVs subtypes contain different HTT protein and RNA levels, important for the development of consistent and reliable biomarkers. Further characterization of neuron-specific EVs content from patient-derived biofluids will lead to the development of novel clinical biomarkers and for evaluation of therapeutic strategies.

## Introduction

Huntington disease (HD) is an inherited autosomal dominant neurodegenerative disorder clinically characterized by progressive motor, cognitive, and psychiatric symptoms [1-3]. HD is caused by an expansion in the cytosine-adenine-guanine (CAG) expansion in the HTT gene that codes for an abnormal polyglutamine tract in the huntingtin protein (HTT) [4]. The polyglutamine-expanded mutant huntingtin (mHTT) leads to the progressive loss of neuronal populations in the striatum, as well as basal ganglia and cerebral cortex [5-10]. Current preclinical studies focus on HTT lowering strategies [11-14]. However, to precisely evaluate these strategies it is essential to identify sensitive and robust biomarkers derived from the brain that are easily accessible in human samples.

Several brain-derived molecules are enriched in the cerebrospinal fluid (CSF) and present biomarkers for HD [15]. Currently, quantification of soluble mHTT in CSF is known to correlate with disease stage and symptoms severity [16, 17]. The development of future assays for HTT measurement from other biofluids, such as blood, will facilitate the access to human samples since CSF collection is an invasive outpatient procedure. Furthermore, the absence of more accessible biomarkers represents an obstacle in the design of future clinical trials of potential therapies. Despite some circulating free HTT in the blood originating from the brain [18], it does not correlate with CSF mHTT or symptom severity [17], possibly because it originates from other sources or is degraded by autoantibodies present at varying levels in circulation.

Another possibility is the isolation of brain-derived extracellular vesicles (EVs) that are detectable in peripheral blood and may reflect the central processes occurring in the brain. EVs are essential vehicles in intercellular communication and mediate various signalling events, carrying cell-specific cargos, such as lipids, proteins, DNA, and RNA [19]. EVs contain surface markers and biological cargo specific to their tissue of origin, justifying their importance not only in normal biology but also in disease as they may report on pathological alterations [20, 21]. EVs is a generic term which includes all types of shedded vesicles, and sizes and mechanisms of biogenesis are the conventional classification approaches for exosomes and ectosomes (also known as microvesicles) [22, 23]. Exosomes are derived from endosomes originating from the multivesicular bodies (MVBs) that are then released upon the fusion of MVBs with the plasma membrane. In contrast, ectosomes (also termed microvesicles) are larger EVs generated by direct outward budding from the plasma membrane.

The incorporation of HTT protein into EVs is consistent with its association with vesicle membranes [24] and role in trafficking secreted proteins [25, 26]. Previous publications have identified EVs as one of several cellular pathways responsible for HTT export from cells to the extracellular space, with particular focus on exosomes [27-32]. It was also shown that CAG-repeat RNA is present in EVs isolated from a cell culture model expressing HTT [28]. However, it is unknown which vesicle type correlates better with HD and focusing only on one subtype restricts our knowledge regarding vesicle biomarkers. In this context, our previous work demonstrated the incorporation of HTT in higher levels in ectosomes compared to exosomes, and neuronal cells exhibited bursting irregularities after their internalization [33, 34]. These results indicate that ectosomes might be a potentially more informative biomarker for HD.

A limitation of biomarker studies in peripheral blood has been the inability to link biomarker levels to brain pathology, as well as the relative uncertainty of their tissue of origin. EVs are released by nearly all cell types in the body, limiting the utility of bulk EV analysis from blood due to contributions from non-central nervous system (CNS) sources [35, 36]. Cells in the CNS secrete EVs to the extracellular space that cross the blood-brain barrier and circulate in CSF and blood [37, 38]. Recent studies have identified markers for the enrichment of CNS cell-type specific EVs from biofluids, allowing for the isolation of neuron-specific EVs from blood [39, 40]. Therefore, development of biomarker approaches using plasma EVs enriched for neural origin has the potential of gaining direct access to brain pathogenic processes. This is relevant to HD, as the selective degeneration and loss of specific neuronal populations, particularly in the striatum, are central to the development of the disease’s motor, cognitive, and psychiatric symptoms [1-3].

Our work investigates the potential of neuronal EVs as non-invasive sources of clinical biomarkers in liquid biopsies to monitor disease progression and therapeutic intervention. We developed an optimized ultracentrifugation protocol for the purification of EVs from human samples and plasma of a humanized HD mouse model, Hu97/18 [41]. Our results show that EVs isolated from different *in vitro* and *in vivo* HD models share common protein composition, highlighting the consistency of EV protein markers across HD models that enables the use of EVs as initial readouts for testing HTT targeting therapies. Consistent with previous published data in other model systems [34, 42], ectosomes isolated from plasma of the Hu97/18 mouse model contain both wild-type (wt) and mHTT in higher levels than in exosomes. Similar results were observed in media from HD induced pluripotent stem cells (iPSCs)-differentiated neurons, Hu97/18 primary neuronal cultures, and human plasma. Interestingly, ectosomes isolated from human plasma also contain higher levels of HTT transcripts. We further demonstrate that a slow freezing protocol for sample storage that lowers temperature by 1°C/min as opposed to the quick-freezing methods typically used for clinical biofluid storage, results in the detection of higher levels of HTT and EV protein markers. This may be related to bursting of EVs during quick freezing, highlighting the importance of a slow freezing protocol for sample storage to improve EV isolation and analysis.

Since EVs can cross the blood-brain barrier (BBB) and circulate in the bloodstream, we developed a protocol to isolate EVs derived from neurons based on previous work [39]. Investigation of HTT levels in neuron-specific EVs from plasma has higher potential as a disease biomarker since these vesicles reflect HTT levels in the CNS. In our work, we demonstrate the feasibility of the isolation of neuron-derived EVs using ATPase Na+/K+ transporting subunit alpha 3 (ATP1A3) immunoprecipitation (IP) from enriched ectosome and exosome fractions [39]. Interestingly, neuron-derived ectosomes also show higher levels of HTT compared with exosomes by immunoblot.

Our results include the development of novel methodology to separate diverse brain-derived EVs from humanized cell models and human biofluid samples, and their biochemical and functional characterization to identify improved EV clinical biomarkers. Importantly, we can isolate neuron-derived EVs from human and mouse plasma and measure HTT levels. Additionally, we show feasibility of isolating EVs from mouse circulation that were produced by striatal neurons. These results reveal the importance of studying brain-specific EVs from patient-derived biofluids and characterization of their contents will be important for the development of clinical biomarkers and therapeutic strategies. The development of assays for CNS HTT protein and transcript measurement from blood will facilitate the access to human samples since CSF collection is invasive. This work will contribute to advancing novel non-invasive clinical biomarkers in liquid biopsies that will allow longitudinal sampling to follow disease progression and evaluate clinical efficacy outcomes.

## Results

### Slow freezing is required to maximally preserve vesicles

Currently, there are no universally implemented processing and storage conditions for EV biomarkers. Poor sample handling and preservation practices may lead to degradation of EVs and thereby compromise the quality of the data obtained. Thus, addressing the impact of processing and storage practices on EVs and their cargo is essential to their use as reliable biomarkers. In our work, we tested how freezing conditions influences HTT levels and EV protein markers in EVs by quick freezing cell media from mouse primary neurons and plasma either by snap-freezing in liquid nitrogen (LN) or placing directly into the freezer (-80°C), which are methods commonly used for storage of clinical biofluids and biobanking. EVs recovered after these quick-freezing methods were compared to slow freezing using a freezing container that decreases temperature slowly by 1°C per minute, a procedure commonly used for storage of live cells to prevent bursting of membranes.

We cultured primary neurons from Hu97/18 model mice [41] and collected media at DIV21 after neuronal differentiation. Media was stored using four different freezing protocols: Freezing container, LN, -80°C, or -80°C with DMSO cryoprotectant, followed by thaw for EV isolation using our optimized ultracentrifugation protocol as previously described [33, 34]. EV characterization was performed in accordance with the MISEV 2018 (minimal information for studies of extracellular vesicles) guidelines [22] and demonstrated in our previous work [33, 34]. Cell lysates and vesicle fractions were assessed by immunoblot for HTT and EV markers, and protein levels normalized to total protein using Memcode (as ectosomes and exosomes do not share a loading control protein at similar levels that can be used for normalization) (**Figure 1**).

**Figure 1.**
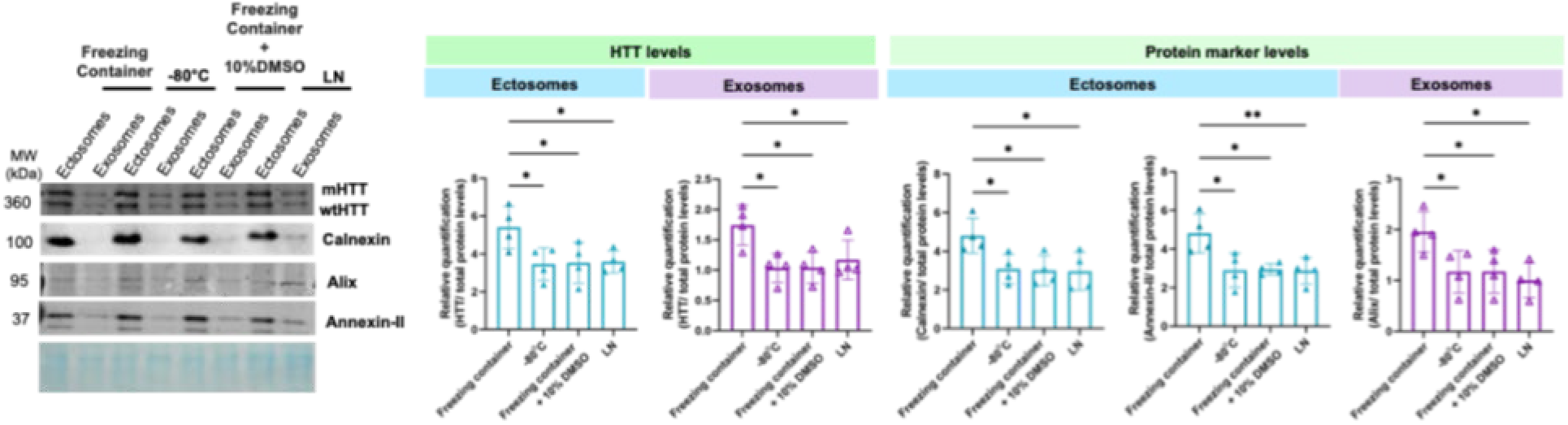
Slow freezing is required to maximally preserve EVs protein markers in cell media collected from Hu97/18 primary neuronal cultures. Cell media was collected at DIV21 from mouse primary neurons expressing wt and mHTT (Hu97/18) and subsequently frozen in different conditions [slow freezing, -80°C, 10% DMSO and liquid nitrogen (LN)]. Immunoblots of ectosomes and exosomes displaying the presence of wt and mHTT. EVs contain specific protein markers, with ectosomes displaying an enrichment of calnexin and annexin-II proteins, while exosomes contain higher levels of alix. Membranes were incubated with the indicated antibodies and protein levels were normalized to total protein levels using MemCode staining (in blue). Immunoblots were cropped for space purposes (N=4). Mean ± SD, *=p<0.05, **=p<0.01. One-way ANOVA with Tukey’s multiple comparisons test.

Our results demonstrate that different freezing conditions affect vesicle properties, and that a slow freezing protocol of cell media results in an improved recovery of ectosomes and exosomes with consequent higher levels of protein markers (**Figure 1**). HTT levels and EV protein markers levels (alix, annexin-II and calnexin) are found at higher levels in vesicles isolated from media frozen with slow freezing protocol (**Figure 1**). Importantly, this result is not dependent on the vesicle size or sample volume loaded into the gel since MemCode is used to normalize protein levels (MemCode staining, in blue in **Figure 1**, shows the total protein levels present in each EV or lysate fraction loaded into the gel). No significant differences were observed between using a cryoprotectant agents (DMSO), -80°C or snap-freezing in liquid nitrogen. This result demonstrates that the slow freezing procedure commonly used for preservation of live cell membranes also maximally preserves EVs and EV HTT.

Furthermore, we collected plasma from Hu97/18 mice to evaluate the effect of freezing conditions on EVs in plasma. We optimized the ultracentrifugation protocol to isolate ectosomes and exosomes from individual animals without pooling plasma samples. Our results demonstrate that slow freezing of mouse plasma also improves EV recovery (**Figure 2**). HTT levels in EVs and protein marker levels (alix and annexin-II) are found at higher levels in vesicles isolated from slow frozen media (**Figure 2**).

**Figure 2.**
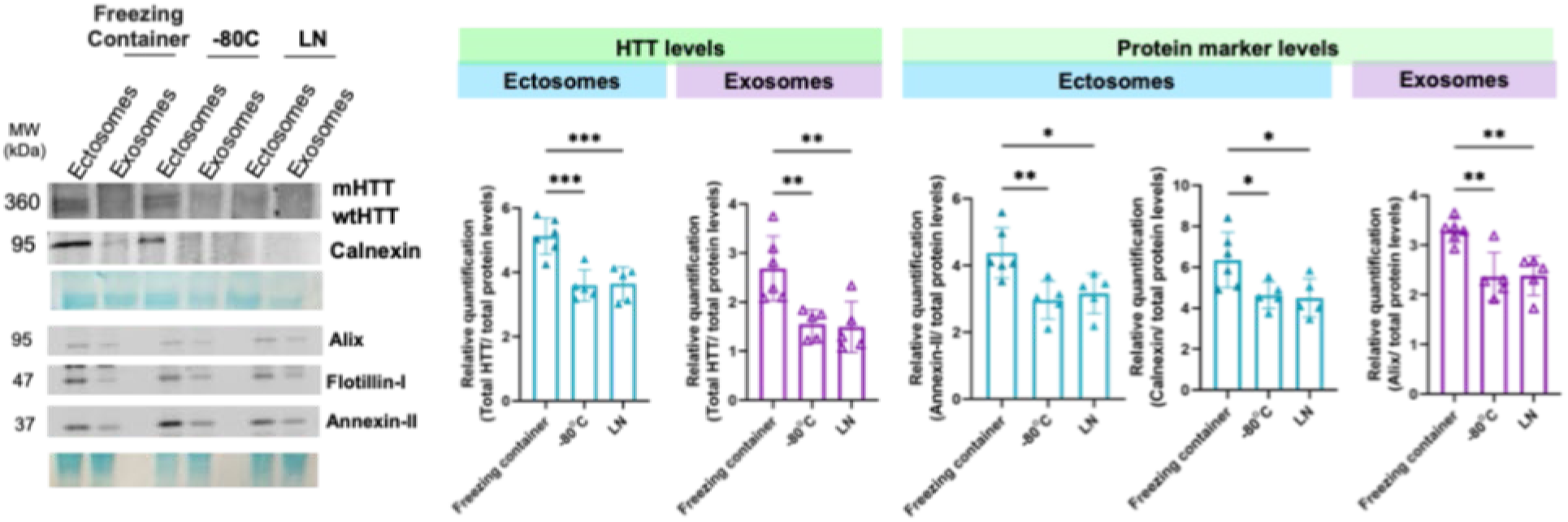
Slow freezing is required to maximally preserve EVs protein markers in plasma collected from Hu97/18 HD mouse model. Plasma was frozen in different conditions [slow freezing, -80°C, and liquid nitrogen (LN)]. Immunoblots of ectosomes and exosomes displaying the presence of wt and mHTT. Membranes were incubated with the indicated antibodies and protein levels were normalized to total protein levels using MemCode staining (in blue). Immunoblots were cropped for space purposes (N=5-6). Mean ± SD, *=p<0.05, **=p<0.01, ***=p<0.001. One-way ANOVA with Tukey’s multiple comparisons test.

These results demonstrate that slow freezing for sample storage that lowers temperature by 1°C/min results in the detection of higher levels of HTT and EV protein markers, highlighting the importance of a slow freezing protocol for sample storage to improve EV isolation and analysis. Maximal preservation of EVs will be important both for the accuracy of biomarker studies as well as enabling the use of smaller starting sample volumes. For our next experiments, we used slow frozen samples to provide the highest vesicles preservation to detect changes between control and HD.

### HTT is present in higher levels in ectosomes than exosomes

Several studies have been focusing on exosomes as biomarkers, however it is not known which vesicle subtype correlates better with disease progression [27]. To evaluate the release of HTT in EVs to the extracellular space, we prepared primary neuronal cultures (Hu97/18) and cell media was collected at DIV21 for ectosome and exosome isolation using our optimized ultracentrifugation protocol as previously described [33, 34]. Cell lysates and vesicle fractions were assessed by immunoblot, and protein levels normalized to total protein using Memcode (**Figure 3**).

**Figure 3.**
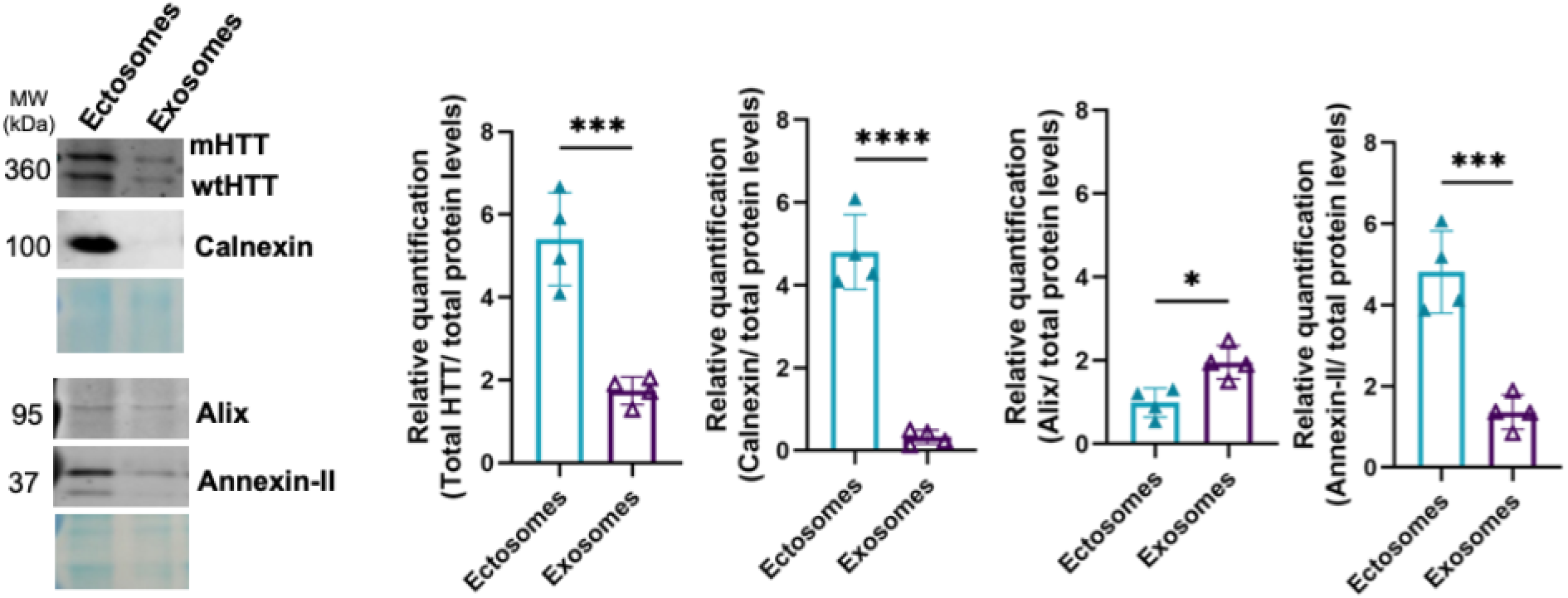
wt and mHTT are secreted in EVs to the extracellular space in mouse primary neuronal cultures. Cell media was collected at DIV21 from mouse primary neurons expressing mHTT (Hu97/18) and subsequently centrifuged at different speeds for EVs isolation. EVs contain specific protein markers, with ectosomes displaying an enrichment of calnexin and annexin-II proteins, while exosomes contain higher levels of alix and flotillin-1. Immunoblots of ectosomes and exosomes displaying the presence of wt and mHTT isolated from EVs. Membranes were incubated with the indicated antibodies and protein levels were normalized to total protein levels using MemCode staining (in blue). Immunoblots were cropped for space purposes (N=4). Mean ± SD, *p<0.05, ***p<0.001, ****p<0.0001. Student’s t test.

As expected, ectosomes had a larger diameter than exosomes (**Supplementary figure 1**), with an average diameter of 93 nm. The average diameter of exosomes was found to be around 60 nm. The conventional exosome protein marker alix was enriched in the exosomal fraction, whereas the ectosomal fraction was enriched in annexin-II and calnexin (**Figure 3**). Both wt and mHTT were detected at higher levels in ectosomes than in exosomes (HTT levels were normalized to the total protein levels in the immunoblot using MemCode) (**Figure 3, supplementary figure 2**). Importantly, this result is not dependent on the vesicle size or sample volume loaded into the gel since MemCode is used to normalize protein levels.

To further confirm our previous results, we isolated EVs from HD iPSC-derived neurons to assess HTT levels (**Figure 4**). Neural stem cells (NSCs) were generated from iPSCs lines originally created from human fibroblasts [43, 44]. NSCs containing 33Q and 109Q were cultured and expanded for differentiation into neuronal cultures (**Supplementary figure 3**), following a previously published protocol [43]. At the early neural progenitor stage of differentiation, cells showed robust expression of NSC markers, including Nestin, Sox1, and Pax6, and no expression of the pluripotency marker Oct4 was observed as shown by immunocytochemistry (ICC) (**Supplementary figure 3**) [43]. At DIV21 after starting the differentiation, we used ICC to confirm that NSCs were differentiated in culture into neurons (**Figure 4**) [43, 45]. Both 33Q and 109Q cultures showed the presence of mature neural markers, including NeuN, MAP2 and β-III tubulin (**Figure 4**). Furthermore, our cultures showed the presence of DARPP32-expressing neurons, a marker for medium spiny neurons (**Figure 4**).

**Figure 4.**
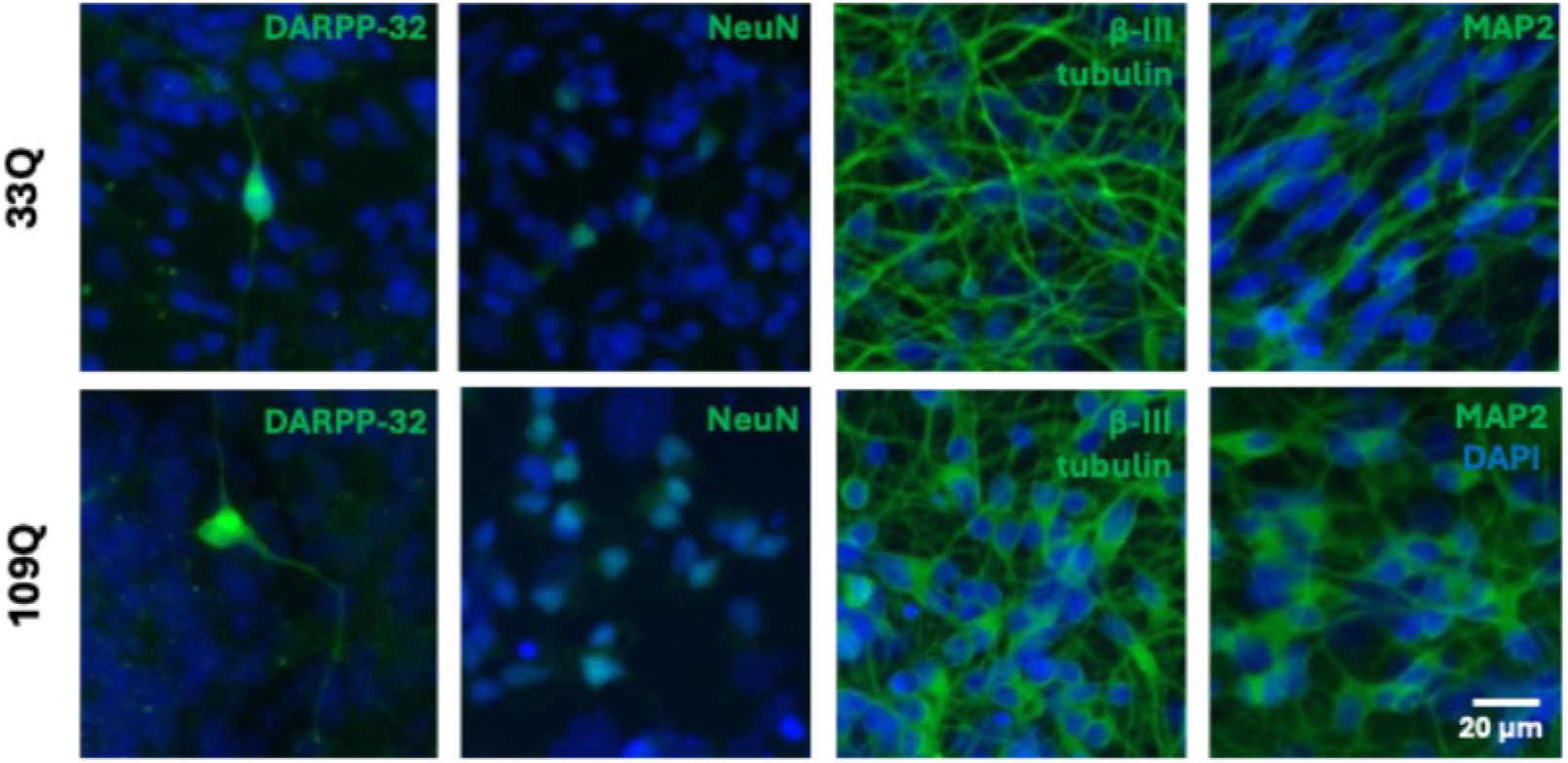
Characterization of iPSC-derived neurons with 33Q and 109Q. Representative images of iPSC-differentiated neurons at DIV21 after starting NSC differentiation protocol until neuronal maturation. Cells were stained for DARPP-32 (green), NeuN (green), β-III-tubulin (green), MAP2 (green) and counterstained with DAPI (blue) to reveal nuclei. Scale bar 20 μm.

We collected cell media from 33Q and 109Q differentiated neurons at DIV21. Our results demonstrate that iPSC-derived neuronal cultures secrete EVs to the extracellular media containing wt and mHTT (**Figure 5**). Vesicle protein markers were enriched as described in our previous results, with alix enriched in exosomes, and flotillin-I, annexin-II and calnexin being in higher levels in ectosomes (**Figure 5**). Immunoblots also showed the expression of wt and mHTT in the protein lysates (**Figure 5**).

**Figure 5.**
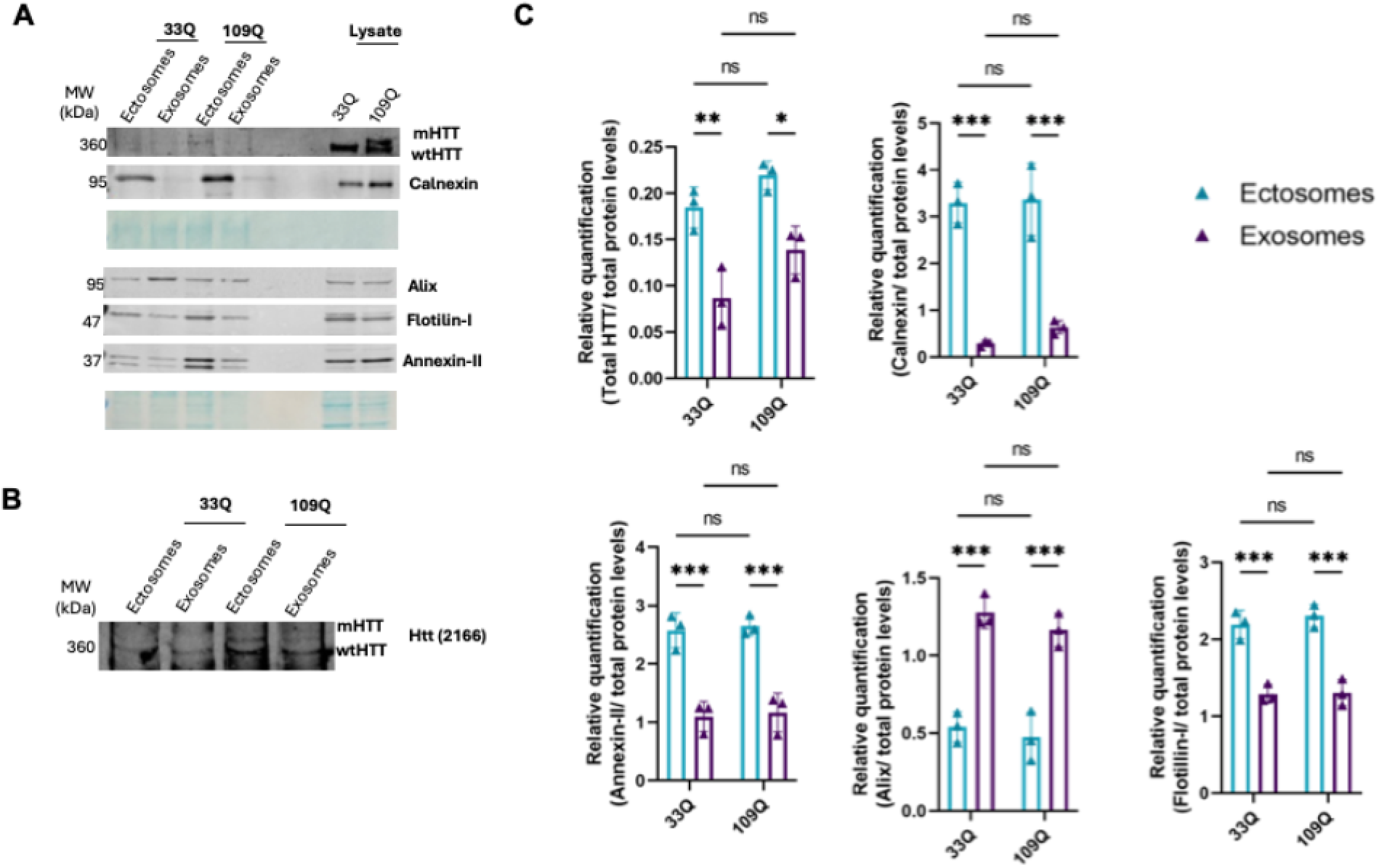
iPSC-derived neuronal cultures from control and HD patients release EVs containing HTT to the extracellular space. Cell media was collected from iPSC-derived primary neurons at DIV21 after starting differentiation and subsequently centrifuged at different speeds for EVs isolation. **(A)** Immunoblots of ectosomal and exosomal fractions purified from the neuronal media. EVs contain specific protein makers, as annexin-II for ectosomes and alix for exosomes. **(B)** Higher magnification and exposition of the HTT bands. **(C)** Quantifications of HTT and EVs protein markers levels. Membranes were incubated with the indicated antibodies and protein levels were normalized to total protein levels using MemCode staining (in blue). Immunoblots were cropped for space purposes (N=3). Mean ± SD, *=p<0.05, **=p<0.01, ***=p<0.001. Two-way ANOVA with Tukey’s multiple comparisons test.

Human iPSC-derived neurons release HTT in higher levels in ectosomes than in exosomes, and this result is not dependent of the quantity of sample loaded into the gel (observed by the same total protein intensities in the MemCode staining).

We also collected plasma from Hu97/18 mice to evaluate the presence of HTT in EVs in circulation. Our results demonstrate that HTT is present in higher levels in ectosomes, however EVs contain low levels of HTT when compared with EVs isolated from neuronal cultures (**Figure 6**).

**Figure 6.**
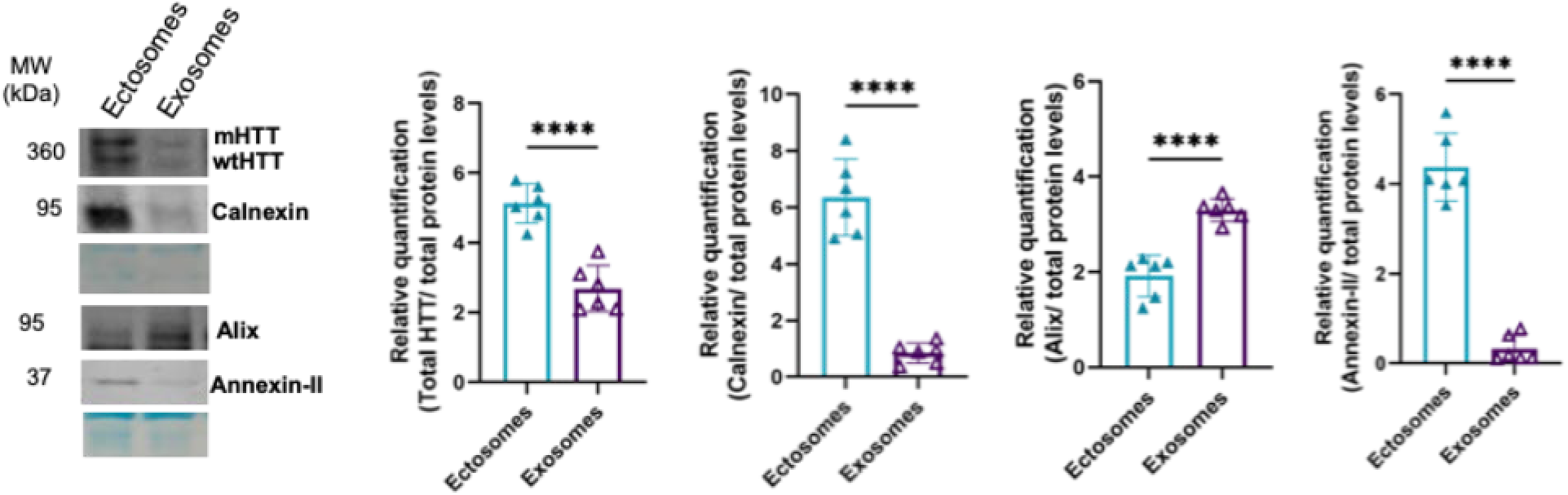
wt and mHTT are present in EVs in circulation in Hu97/18 mouse model. Plasma was collected and subsequently centrifuged at different speeds for EVs isolation. Immunoblots of ectosomes and exosomes displaying the presence of wt and mHTT isolated from Hu97/18 mouse plasma. EVs contain specific protein markers, with ectosomes displaying an enrichment of calnexin and annexin-II proteins, while exosomes contain higher levels of alix. Membranes were incubated with the indicated antibodies and protein levels were normalized to total protein levels using MemCode staining (in blue). Immunoblots were cropped for space purposes (N=6). Mean ± SD, ****p<0.0001. Student’s t test.

Overall, this data supports previous results that indicates both wt and mHTT are incorporated into EVs and released to the extracellular space [33, 34]. EVs showed the same protein markers in EVs isolated from mouse primary neuronal cultures, iPSC-derived neurons and mouse plasma, demonstrating the consistency of EV protein markers across cell models and facilitates future studies of EVs as biomarkers in mouse models.

These results highlight the prominent role that ectosomes, and not only exosomes, may play in the release of HTT to the extracellular space. These results also suggest that HTT may be transported in different EV subtypes in biofluids and emphasize EVs as potential biomarkers in HD.

### Human plasma contains ectosomes and exosomes with HTT

Because EVs can cross the BBB, circulating through the bloodstream and reflecting the cell of origin in terms of disease prognosis and severity, the contents of plasma EVs provide non-invasive biomarkers for HD. Since EVs originating from the brain will express the same surface markers when present in plasma, we evaluated the presence of specific EV protein markers in human plasma and CSF provided by the BioSEND repository (**Figure 7**).

**Figure 7.**
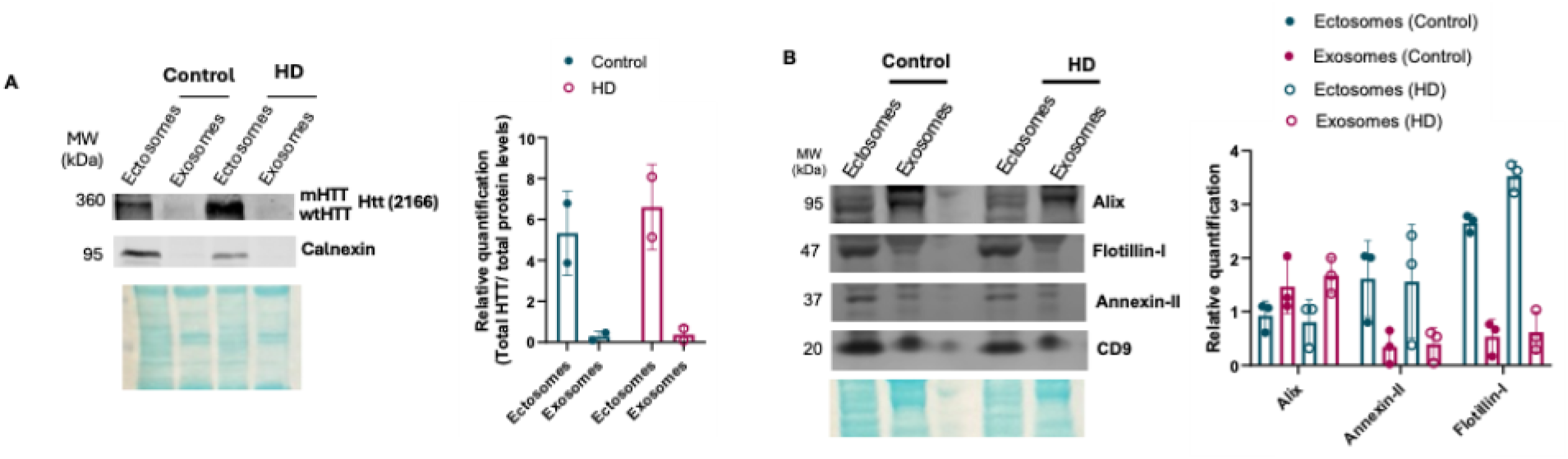
wt and mHTT are present in EVs in circulation in human plasma from BioSEND repository. Plasma was centrifuged at different speeds for EVs isolation. **(A)** Immunoblots of ectosomes and exosomes displaying the presence of wt and mHTT. **(B)** EVs contain specific protein markers, with ectosomes displaying an enrichment of annexin-II protein, while exosomes contain higher levels of alix. Membranes were incubated with the indicated antibodies and protein levels were normalized to total protein levels using MemCode staining (in blue). Immunoblots were cropped for space purposes (N=2-3).

EVs were isolated using the previous optimized ultracentrifugation protocol and vesicle protein markers were evaluated by immunoblot. These results demonstrate that our protocol can separate these EVs from human plasma (**Figure 7**). As shown in **Figure 7**, vesicle protein markers, such as alix and annexin-II, are present in EVs isolated from human plasma, consistent with the previous results in mouse plasma and cell media. HTT seems to be found at higher levels in ectosomes than in exosomes, and this result was not dependent of the quantity of sample loaded into the gel (observed by the same total protein intensities in the MemCode staining) (**Figure 7**).

Furthermore, human CSF also contains EVs however we were not able to detect HTT levels by immunoblot in this sample type with less abundant HTT (**Supplementary figure 4**). Our experiments demonstrate that our protocol can separate ectosomes and exosomes present in CSF and these vesicles contain the same protein markers previously described for cell models and mouse plasma. However, due to the low number of EVs that can be isolated from CSF, we are not able to detect HTT by immunoblot. A starting volume of 5mL of pooled CSF was required to be able to detect these vesicle markers by immunoblot. Compared to the 0.5 ml starting volumes used with plasma samples, this demonstrates the low abundance of EVs in CSF.

We further evaluated the incorporation of HTT transcripts into EVs (**Figure 8**). Our findings support the potential of HTT RNA being present at higher levels in ectosomes compared with exosomes.

**Figure 8.**
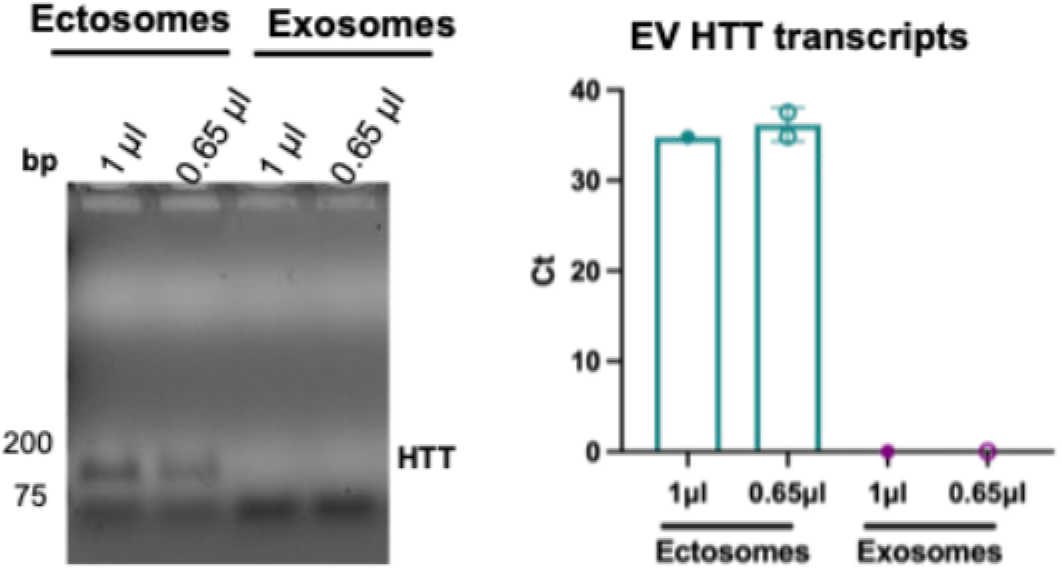
Detection of HTT RNA in EVs isolated from human plasma control sample. Plasma was centrifuged at different speeds for EVs isolation. RNA was extracted from enriched ectosomes and exosomes fractions and loading of HTT RNA into EV was analyses by qPCR. qPCR products of HTT RNA were analyzed on 1.5 % agarose gel. Results are shown in Ct values.

These results indicate that plasma-derived EVs could provide a less-invasive biomarker of brain HTT protein and RNA. The vesicle protein markers are consistent across different cell and animal models, but also in human samples. Furthermore, these results highlight the relevance of ectosomes as a possible biomarker for HD since they contain higher levels of HTT. This result is very important since ectosomes are a less studied vesicle subtype compared with exosomes.

### Isolation of brain-derived EVs using ATP1A3

Currently, the reliability of using L1CAM and other putative neuron-specific proteins as markers of brain neuronal EVs has been questioned [40, 46-48]. Therefore, more reliable and reproducible neuronal markers are needed to isolate and characterize neuronal EVs from human samples. We performed an initial test with L1CAM as a neuronal-specific surface marker in EVs that might be used to enrich for brain EVs in the plasma. Although numerous studies have reported L1CAM-associated biomarker signatures that correlate with disease [49], our preliminary results demonstrate that the available antibodies are unspecific and recognize other proteins present in the plasma (**Supplementary** Figure 5). Furthermore, L1CAM expression is not restricted to neurons and the use of L1CAM alone might result in misleading results (https://www.proteinatlas.org/ENSG00000198910-L1CAM).

A recent study identified adenosine triphosphatase ATP1A3 as a target for isolating neuron-specific EVs from human brain and biofluids in Alzheimer’s disease [39, 40]. ATP1A3 is an integral membrane protein and is mostly enriched in brains with some expression in the heart muscle (https://www.proteinatlas.org/ENSG00000105409-ATP1A3). To confirm the presence of ATP1A3 in our vesicle preparations, we purified ectosomes and exosomes from human and mouse plasma using the optimized ultracentrifugation protocol and analyzed the samples together with human brain lysate (BioSEND fluid samples).

As seen in **Figure 9**, ATP1A3 is present in ectosomes and exosomes from human plasma. Additionally, we used the same protocol to evaluate the presence of ATP1A3 in vesicles isolated from mouse plasma (**Figure 9**). Consistent with our previous results, HTT was present in higher levels in ectosomes than exosomes (**Figure 9**). Our results demonstrate that ectosomes and exosomes contain ATP1A3 and that part of the total plasma vesicle fraction is derived from neurons.

**Figure 9.**
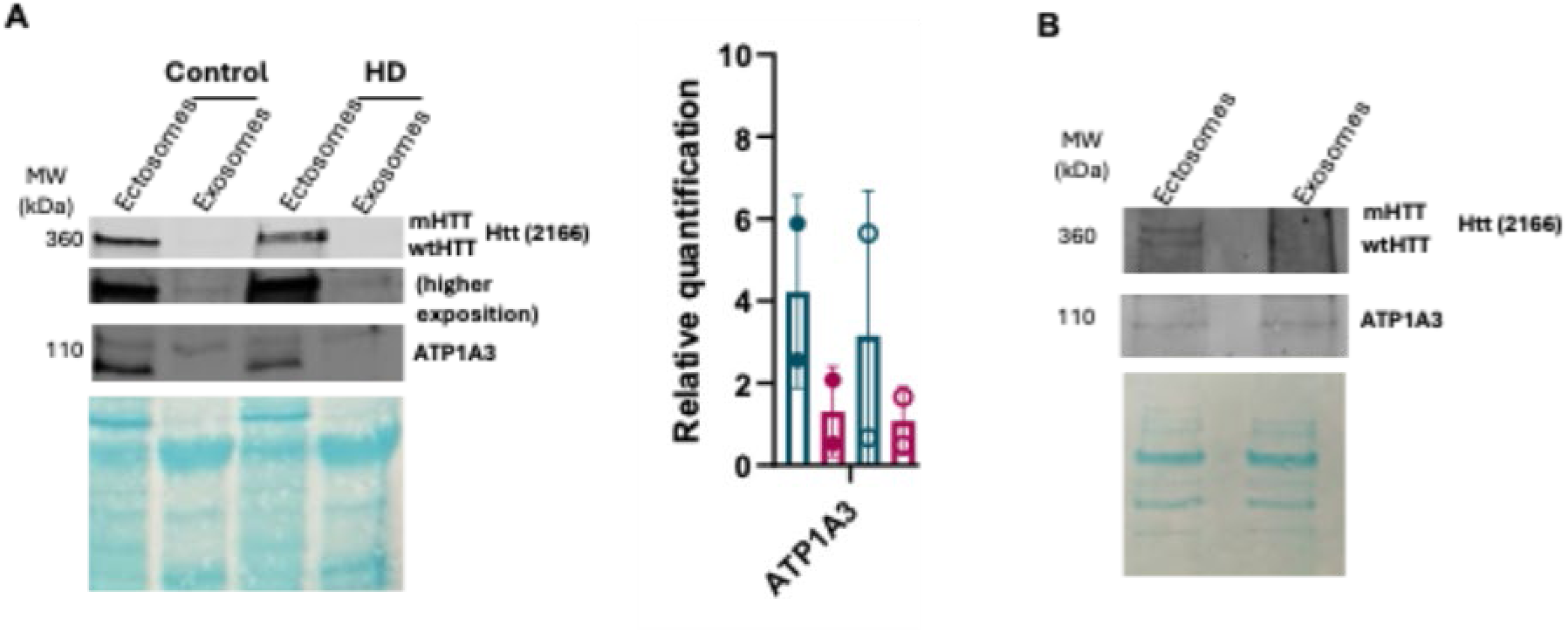
HTT and ATP1A3 are present in EVs isolated from human and mouse plasma. Plasma was centrifuged at different speeds for EVs isolation. **(A)** Ectosomes and exosomes isolated from **(A)** control and HD plasma samples (N=2) and **(B)** mouse plasma (N=1) contain HTT (higher exposition blot shows HTT levels in exosomes) and ATP1A3, a neuron-specific protein. Membranes were incubated with the indicated antibodies and protein levels were normalized to total protein levels using MemCode staining (in blue). Immunoblots were cropped for space purposes.

In our previous work, we determined that expression of GFP in cells results in its incorporation into EVs, providing a useful reporter system for vesicles released from transduced cells in the CNS [42]. To validate the isolation protocol of brain neuron-derived EVs using ATP1A3, we delivered GFP by AAV2 to the striatum of Hu97/18 HD mice to selectively transduce striatal neurons (**Figure 10**). Since EVs can cross the BBB, we expected to find GFP-positive EVs circulating in the plasma. Measurement of GFP levels in EVs was performed using a micro-bead based immunoprecipitation and flow cytometry (IP-FCM) protocol previously described [18]. To validate the GFP IP-FCM protocol, we expressed GFP in HEK cells and isolated ectosomes and exosomes (**Supplementary figure 6**). As previously published, EVs showed the presence of specific protein markers and GFP (**Supplementary figure 6**) [42].

**Figure 10.**
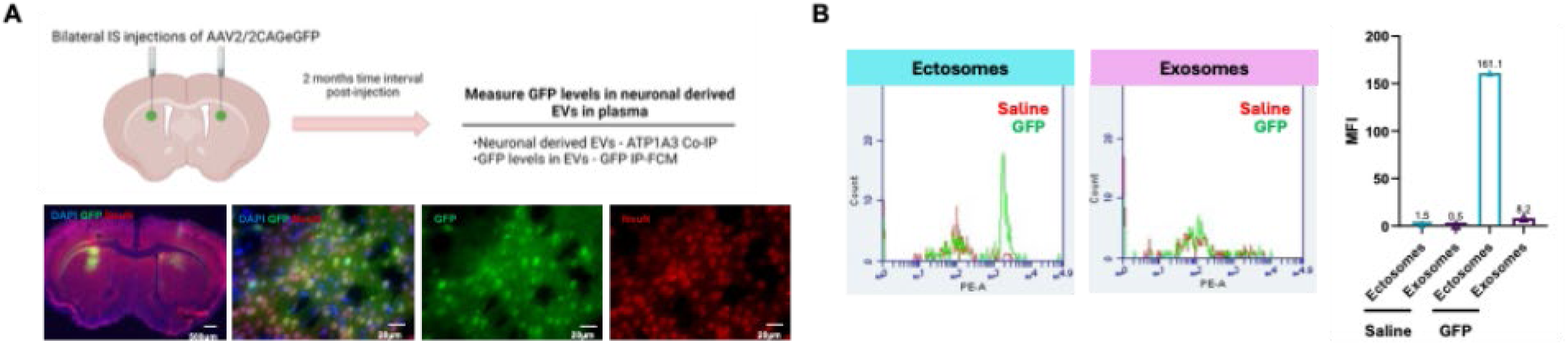
Measurement of GFP signal in brain neuron-EVs present in mouse plasma. 6-month-old Hu97/18 animals received bilateral intrastriatal (IS) injections with AAV2-GFP, and plasma was collected 3 months later for measurement of GFP in EVs. **(A)** Representative images of the distribution of AAV2-GFP in the brain 3 months after intrastriatal (IS) injection. Images show GFP expression (green), neuronal nuclei (NeuN, Red), and all nuclei (DAPI, blue) in the brain (higher magnification of the striatum). **(B)** Quantification of GFP median fluorescence intensity (MFI) in EVs isolated by ATP1A3 IP from plasma EVs using GFP IP-FCM as previously described [18]. EVs from both control (saline) and AAV2-GFP-injected Hu97/18 animals (GFP) were used as input. Beads were also incubated with PBS as a control to evaluate nonspecific binding.

After isolation of ectosomes and exosomes from mouse plasma, we immunocaptured the ATP1A3-positive vesicles by incubating each fraction with magnetic beads coated with ATP1A3 antibody (**Figure 10**) as previously described [39]. Our results demonstrate that neurons release EVs containing GFP to the peripheral blood that can be isolated using ATP1A3 (**Figure 10**).

We have found that ATP1A3 IP captures GFP positive EVs from plasma that was produced in striatal neurons, confirming its utility for isolating CNS neuronal EVs from plasma. In summary, we were able to confirm ATP1A3 as a neuronal marker for the isolation of brain-derived EVs from plasma in HD mice.

### HTT is present in neuron-derived EVs isolated from human plasma

Our previous results show that ATP1A3, a neuron-specific protein, is present in EVs isolated from human and mouse plasma. To validate the isolation protocol of neuron-derived EVs using ATP1A3 and investigate the presence of HTT in these vesicles, control and HD human plasma samples were incubated with magnetic beads coated with ATP1A3 antibody (**Figure 11**). Furthermore, we evaluated the effect of storage conditions in HTT levels in neuronal-derived EVs. For this, human plasma from one control and one HD participant was stored using both slow freezing with a freezing container and fast freezing in a -80°C.

**Figure 11.**
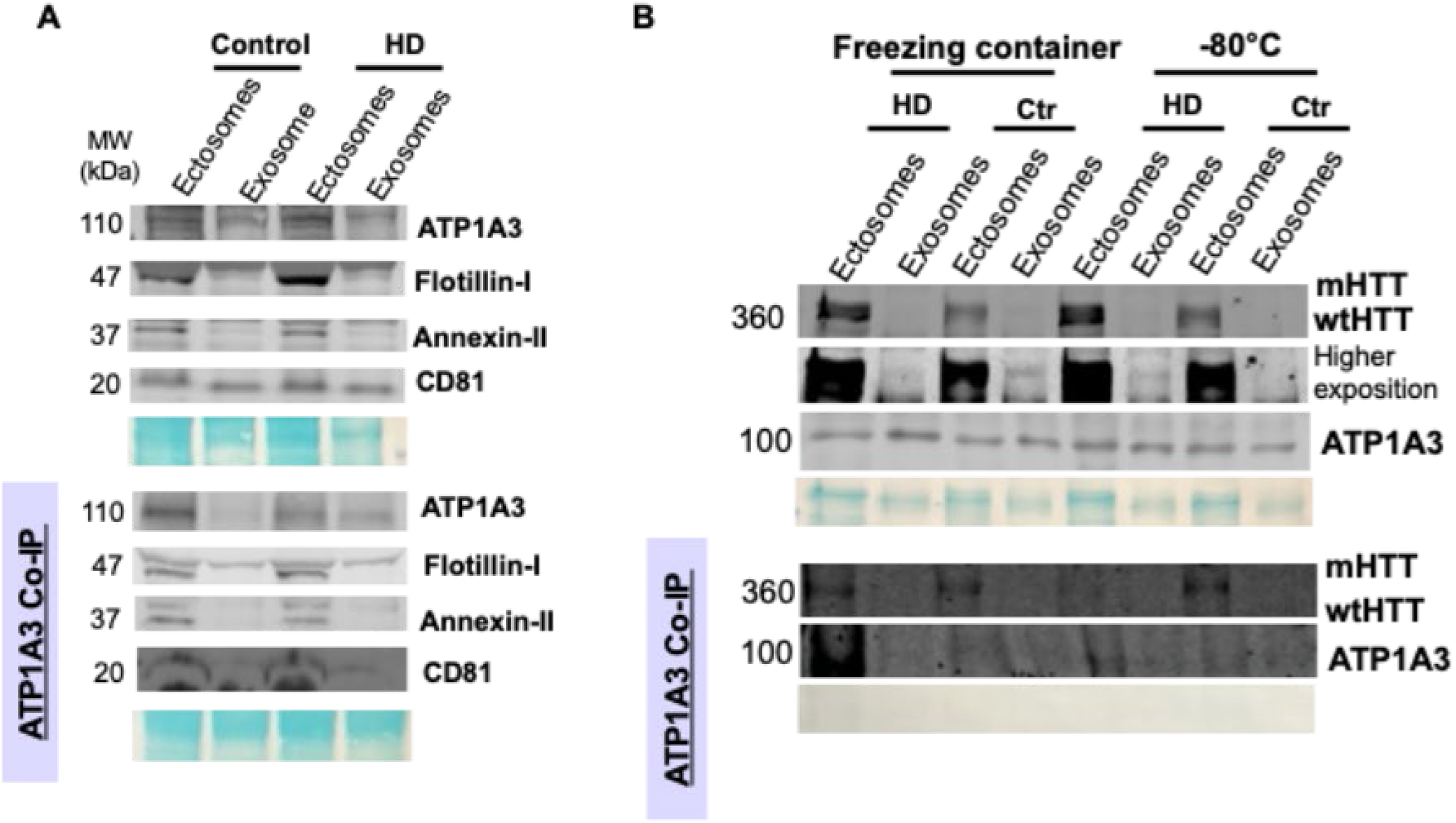
HTT is present in neuron-derived EVs isolated by ATP1A3 IP from human plasma. Enrichment of neuron-derived EVs from human plasma using ATP1A3. Ectosomes and exosomes isolated from control and HD patient plasma were used as input. **(A)** Input and immunocapture results of human samples incubated with magnetic beads coated with ATP1A3 antibody (BioSEND reference pools). Neuronal-derived EVs isolated by IP contain ATP1A3 and specific protein markers (flotillin-I, annexin-II and CD81). **(B)** Human plasma from one control (Ctr) and one HD participant was stored using both slow freezing with a freezing container and fast freezing in a -80°C. Neuronal-derived EVs were incubated with magnetic beads coated with ATP1A3 and contain ATP1A3 and HTT. MemCode staining (in blue) shows the total protein levels present in each EV fraction.

As shown in **Figure 11**, ATP1A3 IP enriched for ectosomes and exosomes containing specific protein markers, as previously described for vesicles isolated from other cell models and mouse plasma. Furthermore, slow freezing improved ATP1A3 and HTT detection, however the levels are very low, making detection difficult by immunoblot (**Figure 11**). These preliminary results demonstrate the feasibility of the isolation of brain-derived EVs using ATP1A3 IP from enriched ectosome and exosome fractions [39]. Furthermore, ectosomes have higher levels of HTT protein compared with exosomes, as observed in our previous results. This result indicates that assessment of HTT protein in EV might be improved by using IP-FCM as is currently used for detection of very low abundance mHTT in CSF [16]. Alternatively, a larger starting plasma sample volume may enable quantification by immunoblot.

Investigation of HTT levels in brain-derived EVs from plasma has higher potential for disease biomarkers since these vesicles reflect HTT levels in the CNS.

## Discussion

Despite the great interest and the increasing number of studies addressing EVs in HD, challenges in EVs isolation, characterization and storage are still open and remain without standardization. The diversity in their size and content suggests that cells may secrete many different types of vesicles, reflecting distinct physiological roles [19]. The use of EVs as biomarkers has attracted significant interest in several areas of research. However, the lack of comprehensive characterization caused an accumulation of contradictory data and challenges in the study of EV biology in neurodegenerative diseases [50]. Furthermore, ectosomes remain largely understudied compared to exosomes [33, 51]. Ectosomes are larger vesicles that bud from the plasma membrane of cells, whereas exosomes originate from multivesicular bodies and are afterwards released to the extracellular space. It is unclear the mechanisms behind inclusion of cellular proteins into EVs, however it appears to be based on controlled protein-sorting mechanisms during their biogenesis, in agreement with unique protein signatures for each EV subtype [23]. Considering the budding biogenesis of ectosomes, they may more faithfully represent cytoplasmic contents than exosomes.

In our work, we developed an optimized ultracentrifugation protocol for the purification of ectosomes and exosomes from cell media, plasma and CSF. Our results show that EVs isolated from different *in vitro* and *in vivo* HD models share common protein composition, highlighting the consistency of EV protein markers across HD models that enables the use of EVs as initial readouts for testing HTT targeting therapies. As previously shown [33], annexin-II is a marker for ectosomes isolated from *in vitro* and *in vivo* HD models but also in human plasma and CSF. Additionally, alix is a marker for exosomes. Calnexin, which is associated with endoplasmic reticulum in the cytoplasm was found to be absent from exosome fractions and detected at very low levels in ectosome fractions in previous studies performed using media from HEK cells [34]. In the more relevant model systems reported here, including cultured neurons, plasma, and brain-derived plasma EV fractions, we find enrichment of calnexin in ectosome fractions. Considering that samples are pre-cleared with centrifugation prior to ultracentrifugation isolation of EVs, we feel it is unlikely that this indicates residual cell or organelle debris, and rather that calnexin is more cytoplasmically available for release in budding ectosomes in these sample types.

We investigated the potential of vesicular HTT, an intracellular protein, that is secreted from cells in EVs. It is currently unclear which EV subtypes correlate with disease progression. Previous studies demonstrated that both wt and mHTT are released from cells in ectosomes and exosomes [28, 29, 32, 33]. Similar to previous work [33], our current findings demonstrate the release of wt and mHTT in higher levels in ectosomes than exosomes using different HD models (mouse primary neuronal cultures, human iPSC-derived neurons, mouse plasma) but also in human samples. It is important to note that these results are not dependent on the vesicle size or sample volume loaded into the gel since MemCode was used to normalize protein levels, since ectosomes and exosomes do not share a loading control protein at similar levels that can be used to normalize the data. Interestingly, we also found higher levels of HTT RNA in ectosomes isolated from control human plasma. This result is very important since it shows that different vesicle types might correlate with disease progression, and ectosomes are possibly a better biomarker for HD since they contain higher levels of HTT protein and transcripts.

While −80°C or LN is a widespread approach for biofluids storage, our study proposes an improved storage strategy for improving the detection of HTT levels in EVs. We demonstrate that a slow freezing protocol for sample storage that lowers temperature by 1°C/min results in the detection of higher levels of HTT and EV protein markers in cell media and human plasma samples, highlighting the importance of a slow freezing protocol for sample storage to improve EV isolation. This result is important for the precise quantification of HTT EVs to correlate with disease progression and testing of HTT lowering therapies. While EV bursting during quick-freezing may not change relationships with disease and only require the use of larger starting sample volumes, we feel it is unlikely that EV bursting would be uniform across different sizes and membrane compositions of EVs, thus quick-freezing may alter disease relationships confounding biomarker data.

A limitation of biomarker studies in peripheral blood has been the inability to connect biomarker levels to brain pathology and the uncertainty of their tissue of origin, that limits the utility of bulk EV analysis due to contributions from non-CNS sources [35, 36]. Previous studies have identified L1CAM and NCAM1 as neuronal markers for the enrichment of CNS cell-type specific EVs from biofluids, however these proteins are not CNS specific and recent studies questioned the reliability of using these proteins as neuron-specific markers of neuronal EVs [40, 46-48]. Recently, ATP1A3 has been described as a neuron-specific marker of brain EVs since its expression is more restricted to neurons in the CNS and enriched in EVs from brain and biofluids [39, 40]. Our work showed the presence of ATP1A3^+^ EVs in mouse and human plasma samples, revealing the access to neuron-derived EVs from a liquid biopsy. Importantly, HTT was present in higher levels in ectosomes, demonstrating the potential of this vesicle subtype as a disease biomarker. We observed a large heterogeneity in HTT levels measured in neuron-derived EVs by IP-FCM. In the future, we will optimize HTT measurement in EVs by IP-FCM to improve the sensitivity and accuracy of HTT detection.

## Conclusions

Our project demonstrates the usefulness of measuring HTT levels in EVs in peripheral blood, giving us a more accessible mechanism for evaluating the success of HTT lowering and HTT targeted clinical trials. Importantly, using slow-freezing, differential ultracentrifugation, and ATP1A3 IP, we demonstrate in mice the successful isolation and detection of GFP protein produced in striatal neurons in neuron-derived ectosomes isolated from blood, validating this EV procedure for biomarkers of brain neurons.

## Materials and methods

### Human plasma and CSF samples

Human samples were collected with an approved protocol, in accordance with the guidelines of the institutional review board, and the full informed consent of the subjects. Plasma and CSF samples were obtained from the BioSEND reference pools or from the Huntington Disease clinic at the University of Central Florida under the Institutional Review Board protocol STUDY00004597. Samples at the Huntington Disease clinic at the University of Central Florida were classified according to the disease diagnostics and information about sex and age were anonymized. Samples were stored in 0.2, mL aliquots at −80°C or using a slow freezing protocol prior to analysis. Samples from the same individual were pooled for EVs isolation from CSF (2mL final volume) and plasma (1mL).

### Mice, treatments, and sample collection

The animal study was reviewed and approved by the Institute Animal Care and Use Committee of the University of Central Florida. Hu97/18 HD model mice [41] were maintained under a 12h light:12h dark cycle in a clean facility and given free access to food and water. Experiments were performed with an approved protocol and in accordance with the guidelines of the animal care committee of the University of Central Florida.

For the AAV-GFP study, virus expressing GFP (AAV2/2CAGeGFP) with 2.02E12vg/ml titer was provided by the Viral Vector Core at the University of Iowa. Bilateral intrastriatal (IS) virus injections with convention enhanced delivery were performed as previously described [52]. Briefly, mice were anesthetized with isoflurane and secured into a stereotaxic frame. The scalp was shaved, sterilized with betaine and 70% ethanol, and an incision made along the midline. The skull was dried to enhance visibility of sutures and landmarks. A dental drill was used to make bilateral burr holes at dorsal-ventral -3.5 to bregma, medio-lateral left +2 and right -2 to bregma, and +0.8 mm anterior-posterior to bregma. A Hamilton syringe with a 30 gauge needle was loaded first with 3 μl of sterile saline and then with 2 μl of either virus diluted in sterile saline. The virus was injected at 0.5 μl/min using an UltraMicroPump with Micro4 controller. The needle was left in place for 5 min and then withdrawn slowly. Plasma and perfused brains were collected 2 months after post-injection for immunoprecipitation followed by flow cytometry (IP-FCM) and immunohistochemical (IHC) analysis, respectively.

For plasma collection, mice were anesthetized using avertin, and whole blood was collected by cardiac puncture and placed into prechilled EDTA-coated tubes (Sarstedt, 41.1395.105) on ice. Whole blood was then centrifuged at 4000 RCF for 10 min at 4°C, plasma was collected and frozen in different conditions until use (flash frozen in liquid N_2_, stored at -80°C, stored using a slow freezing protocol by putting the samples in a freezing container and leave them overnight at−80°C).

For immunohistochemistry (IHC) experiments, mice were perfused transcardially with PBS and 4% PFA. Brains were removed and postfixed in 4% PFA in PBS for 24h at 4°C. The following day, brains were cryoprotected in 30% sucrose with 0.01% sodium azide. Once equilibrated, brains were frozen on dry ice, mounted in Tissue-TEK OCT embedding compound (Sakura, 2580274), and cut via cryostat (Leica Microsystems, CM3050S) into a series of 20 μm coronal sections free-floating in PBS with 0.01% sodium azide.

For evaluation of AVV distribution, a series of sections spaced 200 µm apart and spanning the striatum were stained with GFP antibody (Invitrogen, A-11122, 1:1000) and NeuN (Millipore, MAB377, 1:1000). Primary antibodies were detected with goat anti-rabbit AlexaFluor-488 (Invitrogen, A-11008, 1:500) and goat anti-mouse AlexaFluor-568 (Invitrogen, A-11004, 1:500) secondary antibodies. Sections were then mounted using ProLong Gold Antifade mountant (Invitrogen, P10144) and imaged with either 40x or 63x objectives using a Keyence microscope.

### Neuronal culture

Primary cortical neuronal cultures from Hu97/18 forebrain neurons were prepared as previously described [41]. Hu97/18 transgenic mice at embryonic day (E) E15.5 - E17.5 were removed and stored overnight in Hibernate-E media (Gibco, A1247601) supplemented with 2.5 mL/L GlutaMAX (0.5 mM Gibco, 35050) and 20 mL/L (2%) NeuroCult SM1 Supplement (Stemcell, 05711) during genotyping. In the following day, Hu97/18 cortices with attached striatal tissue were dissected under a stereomicroscope and pooled together. Afterwards, the tissue was transferred to ice-cold 1x Hanks balanced salt solution (HBSS) (Gibco, 24020117) supplemented with 100mL/L D-Glucose (60mg/ml), 10mL/L 1M HEPES Buffer Solution EmbryoMax (Millipore Sigma, TMS-003), and 10mL/L Penicillin Streptomycin 100X Solution (HyClone, SV30010). Tissue was gently dissociated, and cells were resuspended in pre-warmed Neurobasal Medium (Gibco, 21103049) supplemented with 2% NeuroCult SM1 Supplement (Stemcell, 05711), 1% Penicillin/ Streptomycin 100X Solution (HyClone, SV30010) and 0.25% GlutaMAX (0.5 mM Gibco, 35050). Cells were seeded on coverslips coated with 50 µg/mL poly-D-lysine (PDL) (Sigma, P7886) at a density of 250 000 cells per well in 24-well plates and 1x10^6^ cells per well in 6-well plates (Corning) for immunocytochemistry and immunoblot experiments. For immunocytochemistry, coverslips were treated with hydrochloric acid overnight at room temperature (RT) with gentle shaking and washed in ethanol and phosphate-buffered saline (PBS). Coverslips (Marinefield No. 1.5, 13 mm) were then added to wells and dried before PDL coating. Cells were maintained at 37°C with 5% CO^2^/humidified incubator, and fed 1/10^th^ well volume of fresh Neurobasal complete medium twice weekly until day in vitro (DIV) 21.

Media was collected at DIV21 and placed on ice before centrifuging for 10 min at 300x*g* at 4**°**C to pellet cell debris. The supernatant was collected for new tubes and frozen in aliquots using different freezing conditions until use (flash frozen in liquid N2, stored at -80oC, use 10% DMSO, stored using a slow freezing protocol by putting the samples in a freezing container and leave them overnight at −80°C).

### Human HD iPSC-derived neurons differentiation protocol

iPSC generation was previously completed at Cedar’s Sinai Medical Center using non-integrating methods from human fibroblast lines obtained from control and HD patients with CAG repeat sizes of 33 (CS83ictr33n2) and 109 (CS091hd109n4) [43, 45, 53]. Neural stem cells (NSCs) were created using previously published protocols [54, 55]. Coverslips were prepared by washing in hydrochloric acid overnight, followed by washing in ethanol and 1x PBS. Once dry, coverslips were transferred to the bottom of 12-well plates and coated with Matrigel Matrix (Corning, CB-40230). NSCs were cultured onto coverslips and directed toward a striatal fate by plating with neuronal induction media supplemented with GABA (Tocris Bioscience, 0344/1G) and pro-synaptogenic small molecules CHIR99021 (Tocris Bioscience, 44-231-0) and forskolin (Fisher BioReagents, BP25205), as previously published [54-56]. Neurons were directed toward a striatal fate for an additional 9 days, and were kept in culture until DIV21 after starting differentiation upon which neurons were used for immunocytochemistry (ICC) amd media used for EVs isolation.

Media was collected at DIV21 after starting differentiation and placed on ice before centrifuging for 10 min at 300x*g* at 4**°**C to pellet cell debris. The supernatant was collected for new tubes and frozen in aliquots using a slow freezing protocol by putting the samples in a freezing container and leave them overnight at −80°C.

For ICC, fresh 4% PFA (in PBS) was added to the wells and incubated at RT for 15 min. Cells were washed with PBS with 0.1% Tween-20 (PBS-T) and incubated with this solution for 30 min. Afterwards, 5% normal donkey serum (NDS) in PBS-T was added to the wells and incubated at RT for 60 min. Primary antibodies were diluted using PBS-T with 0.2% Bovine Serum Albumin (BSA) and 0.1% sodium azide and incubate overnight at 4°C with slow agitation. NSCs cultures were characterized by ICC using the markers Nestin 10C2 (StemCell, 60091.1, 1:1000), Sox1 JJ20-40 (Thermo Scientific, MA532447, 1:1000) and Pax6 Poly19013 (BioLegend, 901301, 1:1000), and absence of the pluripotency marker Oct4 3A2A20 (StemCell, 60093.1, 1:1000) in the two lines. iPSC-differentiated neurons were stained for DARPP-32 19A3 (Cell signaling, 2306S, 1:1000), NeuN clone A60 (Millipore, mab377), β3-Tubulin D71G9 (Cell signaling, 5568S, 1:1000), MAP2 (Abcam, ab32454, 1:1000) and counterstained with DAPI to reveal nuclei (Thermo Scientific, D1306). Cells were washed with PBS-T and the corresponding secondary antibody was diluted in PBS-T and incubated at RT for 1 hr. Primary antibodies were detected with goat anti-rabbit AlexaFluor-488 (Invitrogen, A-11008, 1:500) and goat anti-mouse AlexaFluor-568 (Invitrogen, A-11004, 1:500) secondary antibodies. Cells were washed with PBS-T and coverslips were mounted using ProLong Gold Antifade mountant (Invitrogen, P10144).

### Human embryonic kidney cells

Human embryonic kidney (HEK) 293 cells (ATTC) were maintained in DMEM medium (Gibco, 11320033) supplemented with 10% FBS (Gibco, 16-000-044) and 1% penicillin-streptomycin (HyClone, SV30010) at 37°C in a 5% CO^2^ atmosphere. Stable cell lines expressing AAV-GFP were developed by viral infection of HEK 293 cells (AAV2/2CAGeGFP provided by the Viral Vector Core at the University of Iowa.). Cells were incubated during 5 days with the virus and after three passages the infection rate was confirmed by microscopy (more than 90% of positive cells).

Media for EVs isolation was collected after 24h and placed on ice before centrifuging for 10 min at 300x*g* at 4**°**C to pellet cell debris. The supernatant was collected for new tubes and frozen in aliquots using a slow freezing protocol by putting the samples in a freezing container and leave them overnight at −80°C.

### Isolation of extracellular vesicles

Isolation of ectosomes and exosomes was performed using an adapted protocol from previous studies [33, 34].

For cell media experiments, the media was collected and placed on ice before centrifuging for 10 min at 300x*g* at 4**°**C to pellet cell debris. The supernatant was collected for new tubes and frozen in aliquots using slow freezing protocol until further analyses. HEK 293 cells were grown in exosome-conditioned medium [depleted of fetal bovine serum (FBS)-derived exosomes], and this medium was produced as previously described [57].

To isolate EVs from cell media, the supernatant was thawed in ice and centrifuged 20min at 2 000x*g*, 4**°**C. All the isolation protocol was performed at 4**°**C and the samples were maintained in ice. 12mL of the supernatant was transferred into ultra-clear tubes (Beckman, NC9194790) and centrifuged in a swing rotor (TH-641 Sorvall) during 90min at 20 000x*g*, 4**°**C. 11mL of the medium was carefully transferred into a new centrifuge tube with a sterile pipette and the pellet containing ectosomes was resuspended in ice cold 1x PBS. The medium was again centrifuged in a swing rotor (TH-641 Sorvall) during 90min at 100 000x*g* (4**°**C) to generate exosomes. The supernatant was discarded and the pellet containing exosomes was resuspended in ice cold 1x PBS with protease and phosphatase inhibitors. Both ectosomes and exosomes pellets were again centrifuged at the correspondent velocities in order to concentrate and clean the pellets. Afterwards the supernatant was removed completely by inverting the tubes and the pellets were resuspended in 100µl of ice cold 1x PBS.

For CSF-derived EVs purification, 2mL of pooled CSF were thawed on ice and diluted in 1X PBS with protease and phosphatase inhibitors to a 13mL final volume. CSF samples were centrifuged at 3000g for 20 min at 4°C. The supernatant was transferred into ultra-clear tubes (Beckman, NC9194790) and centrifuged in a swing rotor (TH-641 Sorvall) during 120 min at 20 000x*g*, 4**°**C. 11mL of the supernatant was carefully transferred into a new centrifuge tube with a sterile pipette and the pellet containing ectosomes was resuspended in ice cold 1x PBS. The supernatant was again centrifuged in a swing rotor (TH-641 Sorvall) during 120min at 100 000x*g* (4**°**C) to generate exosomes. The supernatant was discarded and the pellet containing exosomes was resuspended in ice cold 1x PBS with protease and phosphatase inhibitors. Both ectosomes and exosomes pellets were again centrifuged at the correspondent velocities in order to concentrate and clean the pellets. Afterwards the supernatant was removed completely by inverting the tubes and the pellets were resuspended in 100µl of ice cold 1x PBS.

For plasma-derived EVs purification, 500μL of mouse plasma or 1mL of human plasma were thawed on ice and centrifuged at 1500g for 10 min at 4°C. Samples were then diluted in 1X PBS with protease and phosphatase inhibitors to a 13mL final volume. Plasma samples were centrifuged at 3000g for 20 min at 4°C. The supernatant was transferred into ultra-clear tubes (Beckman, NC9194790) and centrifuged in a swing rotor (TH-641 Sorvall) during 120min at 20 000x*g*, 4**°**C. 11mL of the supernatant was carefully transferred into a new centrifuge tube with a sterile pipette and the pellet containing ectosomes was resuspended in ice cold 1x PBS. The supernatant was again centrifuged in a swing rotor (TH-641 Sorvall) during 120min at 100 000x*g* (4**°**C) to generate exosomes. The supernatant was discarded and the pellet containing exosomes was resuspended in ice cold 1x PBS with protease and phosphatase inhibitors. Both ectosomes and exosomes pellets were again centrifuged at the correspondent velocities in order to concentrate and clean the pellets. Afterwards the supernatant was removed completely by inverting the tubes and the pellets were resuspended in 100µl of ice cold 1x PBS. Protein concentrations were determined by the BCA assay (Thermo Scientific, 23227).

### Immunocapture of neuron-derived EVs using ATP1A3

Immunocapture of neuron-derived EVs from plasma using ATP1A3 was performed according to previous publications [39, 58]. The antibody ATP1A3 (Thermo Scientific, MA3915) was conjugated to Dynabeads M-270 Epoxy using the Dynabeads Antibody Coupling Kit (Thermo Scientific, 14311D) following the manufacturer’s instructions. After isolation of ectosomes and exosomes, equal amounts of each fraction of total plasma-EVs were pretreated with 20 μl of human FcR receptor blocking solution on the ice for 15 min (Miltenyi Biotec, 130-059-901). Afterwards, samples were incubated overnight with ATP1A3-conjugated beads (5 μg/mg) for immunoprecipitation in a final volume of 500 μl at 4°C. Neuron-derived ectosomes and exosomes were eluted in the equal volume of cold SDP lysis buffer [50 mM Tris pH 8.0, 150 mM NaCl, 1% Igepal/NP40, 40 mM β-glycerophosphate (solid), 10 mM NaF and protease inhibitors: 50× Roche Complete, 1× NaVan, 1× PMSF, 1× zVAD] for western blotting. For EV particle analysis, IgG elution buffer (Thermo Scientific, 21028T) was used for the elution, following by pH neutralization with 1 M tris buffer.

### qPCR protocol for HTT transcripts

Ectosomes and exosomes fractions were incubated with TRIzol™ Reagent (Invitrogen, 15596026) and RNA was subsequently purified using RNeasy kit (Qiagen, 74104). cDNA was synthesized using SuperScript III First-Strand Synthesis (Invitrogen, 18080051) following the manufacturer’s protocol. Evaluation of HTT transcript levels was carried out using the following set of primers: F-5’ GAAAGTCAGTCCGGGTAGAACTTC and R– 5’ CAGATACCCGCTCCATAGCAA. Quantitative PCR reactions were carried out using SYBR Green qPCR Master Mix (Thermo scientific, A66732S) according to manufacture protocol. Samples were run on QuantStudio 7 Flex instrument following the manufacturer’s instructions. Data are expressed as Ct. The PCR product along with GeneRuler 1 kb Plus DNA Ladder (Thermo scientific, SM1331) were then run on a 1.5 % agarose gel to verify the amplification product size.

### Western Blot

Brain tissues, cells and EVs fractions were lysed using SDP lysis buffer [50 mM Tris pH 8.0, 150 mM NaCl, 1% Igepal/NP40, 40 mM β-glycerophosphate (solid), 10 mM NaF and protease inhibitors: 50× Roche Complete, 1× NaVan, 1× PMSF, 1× zVAD]. Samples were sonicated for 5 min and protein concentration was measured by microBCA assay (Thermo Scientific, 23235). For quantification of HTT levels, equal amount of sample was separated by 8% low-bis acrylamide gels prepared for allelic separation of HTT [16, 59]. For quantification of other protein levels, equal amount of sample was separated by NuPAGE™ Mini Protein Gels, 4–12 Bis-Tris %, with MOPS running buffer (Invitrogen, NP0336BOX). Proteins were then transferred to 0.45 μm nitrocellulose membranes and incubated with MemCode™ Reversible Protein (Thermo Scientific, 24580) for quantification of the total protein levels following the manufacturer’s instructions. Afterwards, membranes were blocked in 5% dry milk powder (Sigma, 1153630500) in PBS for 1 h at RT and incubated overnight at 4°C in primary antibody in 5% BSA in PBS-T [Huntingtin (1:1000, Millipore, MAB2166MI), calnexin (1:1000, Sigma, C4731), ATP1A3 (1:1000, Thermo Scientific, MA3915) Alix E6P9B (Cell Signaling, 1:1000, 92880), Flotillin-1 D2V7J (1:1000, Cell Signaling, 18634S), TSG101 (1:1000, Abcam, ab30871), Annexin A2 D11G2 (1:1000, Cell signaling, 8235), CD9 (1:1000, Abcam, ab307085), L1CAM UJ127 (1:1000, Abcam, ab20148), CD171 eBio5G3 (1:1000, Invitrogen, 14-1719-82), L1CAM (1:1000, LS Bio, LS-C470565-100), CD81 D3N2D (1:1000, Cell signaling, 56039), GFP (1:1000, Invitrogen, A-11122). Proteins were detected with IR dye 800CW goat anti-mouse (1:250, Rockland, 610-131-007) and AlexaFluor 680 goat anti-rabbit (1:250, Molecular Probes, A21076)-labeled secondary antibodies and imaged using the LiCor Odyssey Infrared Imaging System. Densitometry of bands was performed using Image J software (NIH, version 2.16.0).

### GFP IP-FCM assay

An ultrasensitive micro-bead based immunoprecipitation and flow cytometry (IP-FCM) assay was used to measure GFP levels in EVs as previously described [18]. Briefly, caboxylate-modified latex beads (Invitrogen, C37255) were coupled with the capture antibody GFP (Invitrogen, A-11122) in NP40 lysis buffer (150 mM NaCl, 50 mM Tris, pH 7.4), Halt phosphatase (Thermo Scientific, 78420), and Halt protease inhibitor cocktails (Thermo Scientific, 78429), 2 mM sodium orthovanadate, 10 mM NaF, 10 mM iodoacetamide, and 1% NP40. Capture GFP-coupled beads were then combined with ectosomes and exosomes fractions in triplicate, in a 96-well V-bottom plate (Thermo Scientific, 249944), brought to a total volume of 50 µl in NP40 lysis buffer, mixed well, and incubated overnight at 4°C. On the next day, the plate was spun down for 1 min at 650 RCF and supernatant was removed. Beads were washed with IP-FCM wash buffer (100 mM NaCl, 50 mM Tris, pH 7.4, 1% BSA, 0.01% sodium azide). GFP antibody (Invitrogen, A-11122) was biotinylated using EZ-Link Sulfo-NHS-Biotin (Thermo Scientific, 21217), and 50 µl of the diluted antibody was added to each well plate and incubated for 2 h at 4°C. Beads were washed with IP-FCM wash buffer and 1:200 Streptavidin-PE (BD Biosciences, 554061) was added to each well and incubated at RT protected from light for 30 min. Beads were washed and resuspended with IP-FCM buffer, and fluorescence intensity was measured using an Acuri C6 flow cytometer (BD Biosciences). Median fluorescence intensity (MFI) of PE for GFP was measured for each sample to determine target protein levels. MFI of PE was corrected for background signal and presented as absolute MFI values.

### NTA analysis

Particle number and size distribution in ectosomes and exosomes samples were determined by ZetaView tracking analysis (Particle Metrix). The system was calibrated using 100 μm polystyrene beads (Applied Microspheres, 700074). Samples were diluted in 1x PBS to ∼10^7^ particles/ml to a final volume of 5mL prior to analysis, according to the manufacturer recommendations. ZetaView software (version 8.02.28) was used for the analysis of light scattering at 11 camera positions with 2 s video lengths, a camera frame rate of 15 frames per second, with a system temperature of ∼21 °C to obtain the size profiling and quantification of isolated EVs.

### Imaging

Imaging was performed on Keyence microscope using the company software with 10-40x objective lenses. The acquisition settings were optimized to avoid underexposure and oversaturation effects and kept equal throughout image acquisition of the samples.

### Quantification and statistical analysis

Images were analyzed using ImageJ software (NIH, version 2.16.0) [60]. All data are presented as mean ± SD. Data from at least three independent experiments and each replicate represents one independent experiment. To assess differences between two groups, two-tailed unpaired student t test was performed using GraphPad Prism 10 software (GraphPad). To assess differences between more than two groups, significant differences were assessed by one-way ANOVA followed by multiple comparisons with significance between groups corrected by Tukey’s multiple comparisons test using GraphPad Prism 10 software (GraphPad). Differences were considered to be significant for values of p < 0.05 and are expressed as mean ± SD.

## Author contributions

A.L.S. and I.C.B. conceived the study and designed experiments. I.C.B and Y.X. performed experiments. I.C.B performed data analysis and interpretation. I.C.B generated the graphs and figures. I.C.B. drafted the manuscript. I.C.B. and A.L.S. revised the manuscript. A.L.S. supervised the work. All authors contributed to the article and approved the submitted version.

## Data Availability Statement

The datasets generated for this study are available on request to the corresponding author.

## Acknowledgments

We would like to thank Zena Khaled and Ratnesh Kesineni for assistance with mouse husbandry and genotyping, The BioSend repository for providing pooled HD and control plasma and CSF samples for assay development and characterization, and the University of Iowa Viral Vector Core for AAV production.

