## Supplemental Information for "Neuron-derived extracellular vesicles in plasma present a potential non-invasive biomarker for Huntingtin protein and RNA assessment in Huntington disease"

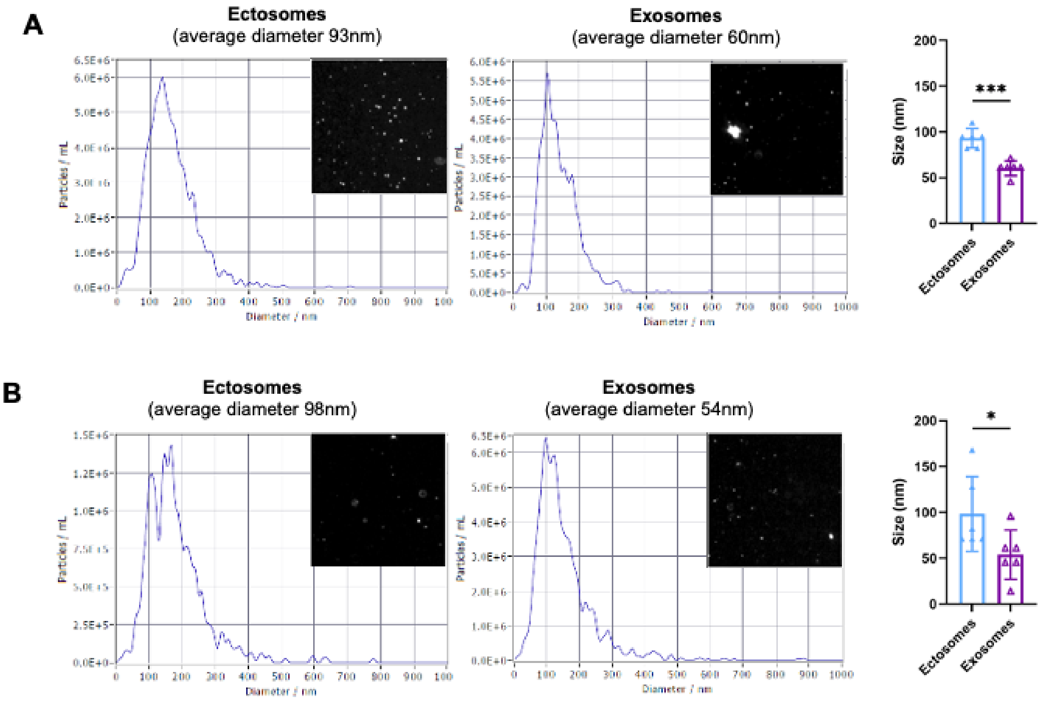


**Supplementary figure 1. Ectosomes exhibit higher average size compared with exosomes.** Representative size distribution by intensity measured by ZetaView of **(A)** EVs isolated from primary neurons cell media and **(B)** isolated from mouse plasma. Average size distribution of ectosomes and exosomes is shown on the right. Mean ± SD, *p<0.05, *** =p<0.001. Student’s t test.

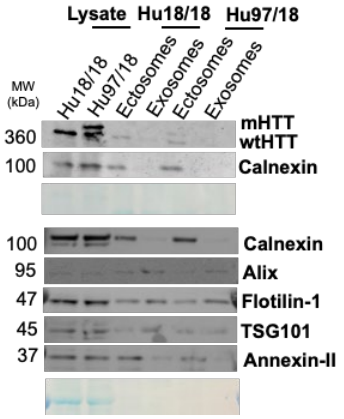


**Supplementary figure 2**. **wt and mHTT are secreted in EVs to the extracellular space in mouse primary neuronal cultures.** Cell media was collected at DIV21 from mouse primary neurons expressing wt (Hu18/18) or mHTT (Hu97/18) and subsequently centrifuged at different speeds for EVs isolation. EVs contain specific protein markers, with ectosomes displaying an enrichment of calnexin and annexin-II proteins, while exosomes contain higher levels of alix and flotillin-1. Membranes were incubated with the indicated antibodies and protein levels were normalized to total protein levels using MemCode staining (in blue). Immunoblots were cropped for space purposes.

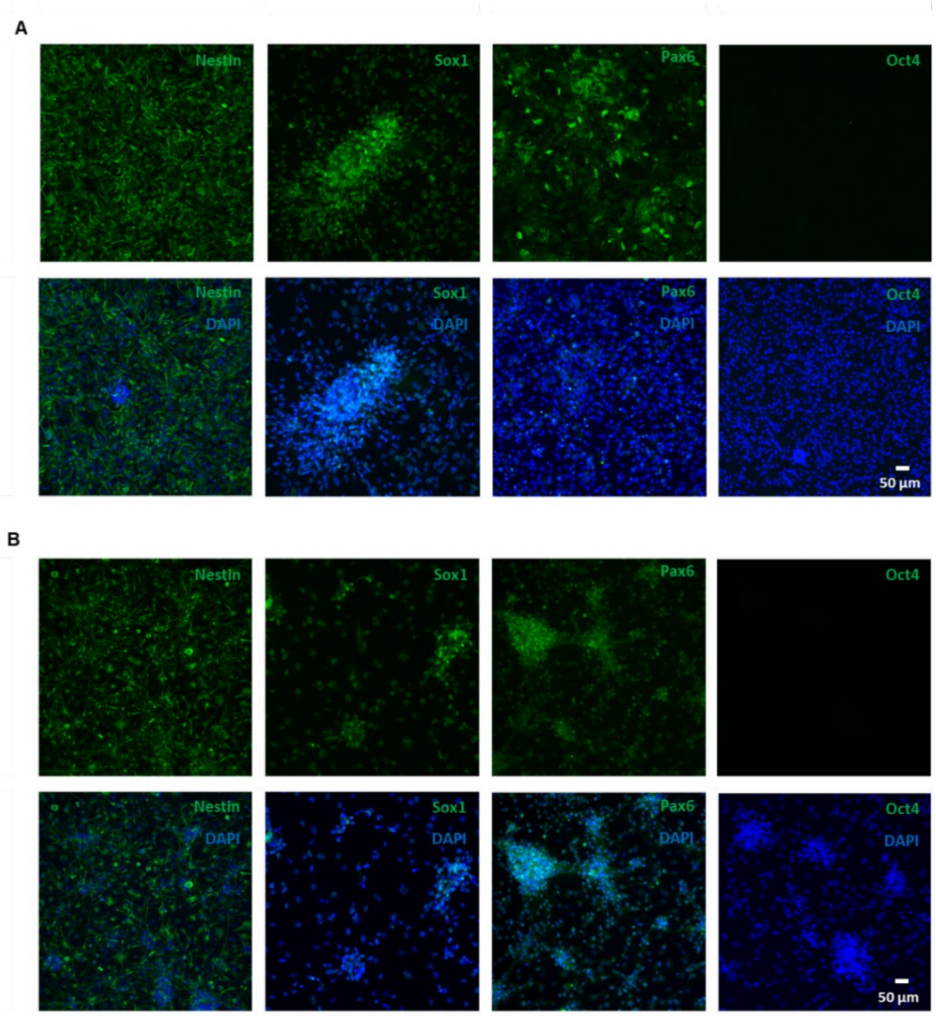


**Supplementary figure 3. Characterization of HD NSCs lines with 33 and 109 polyglutamines.** NSCs with **(A)** 33Q or **(B)** 109Q repeats were expanded as spherical aggregates in a self-renewing condition. ICC demonstrates the expression of the neural progenitor markers Nestin, Sox1 and Pax6, and absence of the pluripotency marker Oct4 in the two lines. Bottom figures show cells counterstained with DAPI to reveal nuclei.

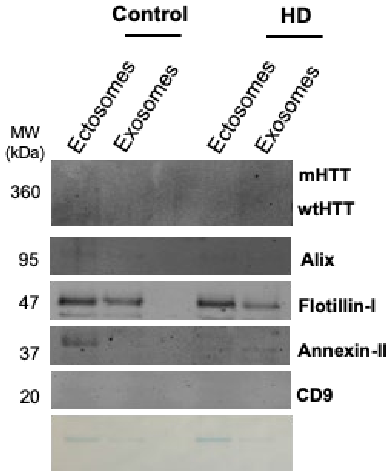


**Supplementary figure 4. Ectosomes and exosomes are present in human CSF from BioSEND reference pools).** Immunoblots of ectosomal and exosomal fractions purified from Human CSF. EVs contain specific protein makers, as annexin-II for ectosomes and alix for exosomes. Membranes were incubated with the indicated antibodies and total protein levels are shown using MemCode staining (in blue). Immunoblots were cropped for space purposes (N=1).

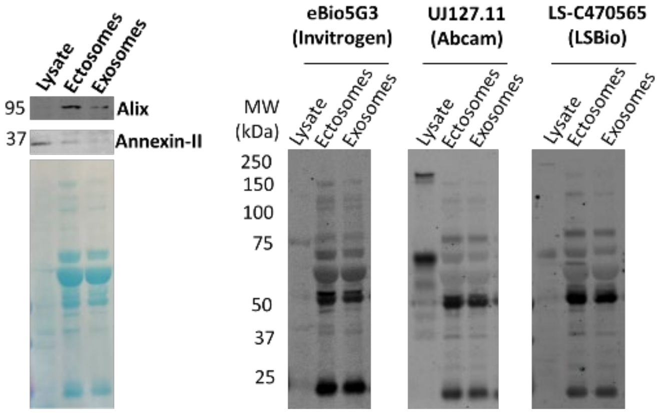


**Supplementary Figure 5. Presence of L1CAM in EVs isolated from human plasma.** EVs isolated from human plasma contain specific protein makers, annexin-II for ectosomes and alix for exosomes. EVs fractions were incubated with different L1CAM antibodies that recognize different regions of the protein. MemCode staining (in blue) shows the total protein levels present in each EVs and human brain lysate fraction.

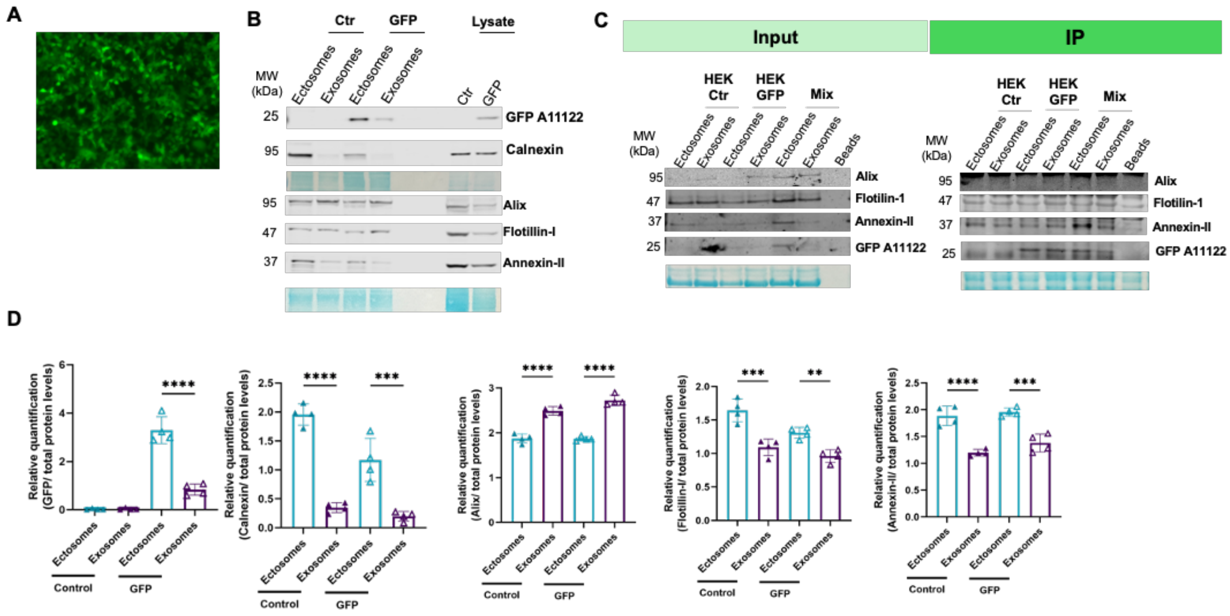


**Supplementary figure 6. HEK cells stably expressing GFP secrete EVs to the extracellular space contain GFP.** Stable cell lines created by infecting HEK cells with AA2-GFP were created. Cell media was collected from cells after 24h and subsequently centrifuged at different speeds for EVs isolation. **(A)** Representative image of the distribution of GFP in HEK cells. **(B)** Immunoblots of ectosomal and exosomal fractions purified from the cell media isolated of control (ctr) and GFP-expressing cells (GFP). EVs contain specific protein makers, as annexin-II for ectosomes and alix for exosomes. **(C)** Enrichment of GFP-positive EVs from HEK cell media. Input and immunocapture results of samples incubated with magnetic beads coated with GFP antibody. Ectosomes and exosomes from control (Ctr) or GFP-expressing cells (GFP) were used as input, as well as a 1:1 mixture of EVs from Ctr and GFP expressing cells (Ctr+GFP). Beads were also incubated with PBS as a control to evaluate nonspecific binding. **(D)** Quantifications of GFP and EVs protein markers levels. Membranes were incubated with the indicated antibodies and protein levels were normalized to total protein levels using MemCode staining (in blue). Immunoblots were cropped for space purposes (N=4). Mean ± SD,$^{**}=p<0.01$, $^{***}=p<0.001$, $^{****}=p<0.0001$. Two-way ANOVA with Tukey's multiple comparisons test.
